# The *β-amylase7* gene in *Zea mays* encodes a protein with structural and catalytic properties similar to Arabidopsis BAM2

**DOI:** 10.1101/2021.10.14.464379

**Authors:** Claire M. Ravenburg, McKayla B. Riney, Jonathan D. Monroe, Christopher E. Berndsen

## Abstract

Starch accumulates in the plastids of green plant tissue during the day to provide carbon for metabolism at night. Starch hydrolysis is catalyzed by members of the β-amylase (BAM) family, which in *Arabidopsis thaliana* (At), includes nine structurally and functionally diverse members. One of these enzymes, AtBAM2, is a plastid-localized enzyme that is unique among characterized β-amylases since it is tetrameric and exhibits sigmoidal kinetics. Sequence alignments show that the BAM domains of AtBAM7, a catalytically inactive, nuclear-localized transcription factor with an N-terminal DNA binding domain, and AtBAM2 are more closely related to each other than they are to any other AtBAM. Since *BAM2* is found in more ancient lineages, it was hypothesized that *BAM7* evolved from *BAM2*. However, analysis of the genomes of 48 flowering plants revealed 12 species that appear to have a *BAM7* gene but lack a *BAM2* gene. Upon closer inspection, these BAM7 proteins have a greater percent identity to AtBAM2 than to AtBAM7, and they share all of the AtBAM2 functional residues that *BAM7* proteins normally lack. We hypothesize that these genes may encode a BAM2-like protein although they are currently annotated as BAM7-like genes. To test this hypothesis, we designed a cDNA of the short form of corn BAM7 (ZmBAM7-S) for expression in *E. coli*. Small Angle X-Ray Scattering data indicate that ZmBAM7-S has a tetrameric solution structure more similar to that of AtBAM2 than AtBAM1. In addition, partially purified ZmBAM7-S is catalytically active and exhibits sigmoidal kinetics. Together these data suggest that some *BAM7* genes may encode a functional BAM2. Exploring and understanding β-amylase gene structure could have impacts on the current annotation of genes.

## Introduction

In most plants, starch provides the carbon and energy necessary to sustain metabolism at night when photosynthesis is inactive or after a long period of dormancy (Zeeman *et al.*, 2010). One group of plant proteins involved in starch metabolism is the β-amylase (BAM) family (Monroe & Storm, 2018; Thalmann *et al.*, 2019). BAM enzymes catalyze the hydrolysis of α-1,4 glycosidic bonds in starch which releases maltose (Zeeman *et al.*, 2010). In *Arabidopsis thaliana* (At), there are nine members of the BAM family all of which are encoded by separate genes, have conserved BAM domains, and include an N-terminal variable region involved in localization of the proteins (Monroe & Storm, 2018; Thalmann *et al.*, 2019).

Five of the Arabidopsis BAMs are catalytically active on starch or dextrin products (BAM1, BAM2, BAM3, BAM5, and BAM6) while the other four are non-catalytic on these substrates (BAM4, BAM7, BAM8 and BAM9) (Monroe & Storm, 2018). The majority of the BAMs in Arabidopsis are thought to function as monomers while some, such as BAM7 and BAM8, are predicted to form dimer complexes with themselves or each other (Sparla *et al.*, 2006; Reinhold *et al.*, 2011; Soyk *et al.*, 2014; Monroe & Storm, 2018). Additionally, AtBAM2 and *Ipomoea batatas* (Ib) BAM5 were identified as tetramers (Cheong *et al.*, 1995; Monroe *et al.*, 2017, 2018; Chandrasekharan *et al.*, 2020). In addition to their structural and functional differences, the members of the AtBAM family also vary in their cellular localization; only BAM5 is exclusively cytosolic while both BAM7 and BAM8 are nuclear. The remaining six AtBAMs, including BAM2, are found in plastids where starch is stored suggesting that they might be involved in starch metabolism (Monroe & Storm, 2018). Although there has been significant attention dedicated to some BAM proteins, much remains to be understood about the structures, in-vivo functions, and evolutionary relationships of the BAM proteins.

This work focuses on comparing BAM2 and BAM7 proteins from Arabidopsis and *Zea mays* (Zm; corn). Based on conserved intron positions, BAM2 is the proposed ancestral protein of the BAM subfamily 2 which includes BAM4, BAM5, BAM6, BAM7, and BAM8 (Monroe and Storm, 2018). Although we do not yet understand BAM2’s function, it has persisted in nearly all land plants. Additionally, the *BAM7* gene likely arose by the fusion of a gene encoding a BZR1-like DNA-binding domain to the 5’ end of *BAM2* (Reinhold *et al.*, 2011; Soyk *et al.*, 2014; Thalmann *et al.*, 2019). However, the sequences of BAM2 and BAM7 across land plants have significant differences within the catalytic residues of their respective BAM domains (Monroe *et al.*, 2017). While all annotated *BAM2* genes encode the residues necessary for catalytic activity, most *BAM7* genes have mutations in at least 3 of the 15 catalytic residues, and this likely contributes to BAM7 proteins being catalytically inactive (Reinhold *et al.*, 2011; Soyk *et al.*, 2014). Conservation of residues and sequence identity is the most reliable method when classifying BAMs (Monroe & Storm, 2018).

In this work, we have used conservation of residues to identify that the *BAM7* gene in some species such as corn may contain two transcriptional start sites that encode two different BAM proteins (ZmBAM7-L and ZmBAM7-S). We further characterized the catalytic activity and solution structure of ZmBAM7-S and compared these findings to AtBAM2, IpBAM5, and BAM1 (Cheong *et al.*, 1995; Chandrasekharan *et al.*, 2020). This work expands on our current understanding of both BAM gene structure and protein form variability. Ultimately, we find that ZmBAM7-S shows sigmoidal saturation kinetics and a tetrameric structure that suggests that it is a BAM2-like β-amylase.

## Materials and Methods

### Protein Sequence Alignments

Using the At*BAM2* sequence as a reference (NP_191958.3), a BLAST search was conducted using the NCBI RefSeq database to identify *BAM2* and *BAM7* genes in other annotated land plant genomes. Similarly, the sequence of At*BAM1* (NP_189034.1) was used to find *BAM1* genes in land plants to compare to the *BAM2 and BAM7* genes. The FASTA-formatted protein sequences were then downloaded from NCBI (https://www.ncbi.nlm.nih.gov/). These protein sequences were submitted to Clustal Omega (https://www.ebi.ac.uk/Tools/msa/clustalo/) to obtain multiple sequence alignments (Madeira *et al.*, 2019).

### Protein Start Site Predictions

The DNA sequence of *BAM7* genes were manually analyzed for alternative in-frame translational start sites (ATG) in the intron between the BZR1 and BAM domains. After *in silico* translation, cellular locations for predicted full-length BAM7 proteins and shorter proteins lacking the BZR1 domain and starting from in-frame Met residues were conducted using LOCALIZER (Sperschneider *et al.*, 2017)).

### Expression Vector Construction

DNA sequences of At*BAM2* (NP_191958.3) and Zm*BAM7* (NP_001337631.1) were obtained from NCBI (https://www.ncbi.nlm.nih.gov/). The sequence of Zm*BAM7-S* was determined based on its in-frame start codon prediction 42 bases 5’ to the start of exon 2. The lengths of the predicted chloroplast transit peptides were determined using TargetP-2.0 (Armenteros *et al.*, 2019). Synthesis of the At*BAM2* and Zm*BAM7-S* coding sequences lacking the predicted 55 and 66 residue chloroplast transit peptides, respectively, was carried out by GenScript using codon optimization for expression in *E. coli* (sequence available as Supplemental Information). The cDNAs were then cloned into pET-15b such that the expressed proteins would contain an N-terminal His-Tag (GenScript, Piscataway, NJ). Transformation of competent DH5α *E. coli* cells (New England BioLabs, Beverly, MA) with the plasmid DNA was carried out using the manufacturer’s protocol. Plasmids were isolated by miniprep and confirmed after digestion with *Bam*HI and *Nde*I. Transformation of BL21 *E. coli* cells with each plasmid DNA was carried out using the Rapid colony transformation procedure (Micklos & Freyer, 1990). The BAM1 cDNA was a gift from Heike Reinhold and was described previously (Monroe *et al.*, 2014).

### Protein Expression and Purification

Cell cultures of BL21 lacking any plasmid (control) or containing one of the previously described recombinant plasmids were grown to an optical density of 0.7 at 600 nm in Luria-Bertani media with 100 μg mL^−1^ carbenicillin at 37 °C and with shaking at 250 rpm. Isopropyl β-D-1-thiogalactopyranoside was added to a final concentration of 0.8 mM, and the flasks were shaken at 250 rpm at 20 °C overnight. Cells were pelleted by centrifugation at 3,000 rcf for 15 minutes at 4 °C then frozen at −80 °C for at least 10 minutes. Cell pellets were thawed and resuspended in 50 mM NaH_2_PO_4_, 0.5 M NaCl, 0.2 mM tris(2-carboxyethyl)phosphine (TCEP), and 2 mM imidazole (pH 8.0) with EDTA-free protease inhibitor tablets (Pierce A32965). Cell lysis was completed by sonication in an ice bath (Misonix S-4000; Microtip) for 2.5 minutes at 55% amplitude (5 seconds on; 20 seconds off). After centrifugation of the cell lysate at 17,418 rcf for 20 minutes at 4 °C, the supernatants of AtBAM2 and ZmBAM7-S were separately loaded onto a TALON cobalt affinity column using an ÄKTA Start system (Cytiva Life Science, Marlborough, MA). Bound proteins were eluted from the TALON cobalt column using a stepwise addition of a second buffer containing 50 mM NaH_2_PO_4_, 0.5 M NaCl, 0.2 mM TCEP, and 200 mM imidazole (pH 8.0). The four 10 mL elution steps contained 12.5, 50, 125, or 200 mM imidazole mixed by the ÄKTA Start system. Fractions were analyzed for purity by SDS-PAGE using BioRad Mini-PROTEAN TGX Stain-Free gels in a buffer containing 250 mM Tris base, 1.92 M glycine, and 1% w/v SDS. Precision Plus Protein Unstained Standards (BioRad) were used as a marker for protein size. Dialysis using SpectraPor tubing (Spectrum, New Brunswick, NJ) with a molecular weight cutoff of 6-8 kDa was completed overnight at 4 °C with constant stirring in 2 L of a buffer containing 20 mM HEPES, pH 7, 100 mM NaCl, and 0.2 mM TCEP. The dialyzed proteins were concentrated in an Amicon Ultra-15 concentrator with a molecular weight cutoff of 10 kDa at intervals of 30 minutes at 5,000 rcf and 4 °C until the desired volume was reached (~1.2 mL concentrated from 50 mL). The concentration of protein was determined using the Bio-Rad Protein Assay Kit with BSA as the standard.

The plasmid of AtBAM1 was added to BL21 cell cultures. The cells containing the plasmid for AtBAM1 were then grown in 2xYT media (RPI) along with 1 μL mL^−1^ Kanamycin at 37 °C with shaking at 200 rpm. Once the growth reached an optical density of 0.6 at 600 nm, 0.75 mM of IPTG was added and the temperature was dropped to 20 °C. Cells shook overnight for about 10-12 hours, and then were pelleted by centrifugation at 5,000 rpm (3,024 rcf) at 4 °C for 15 minutes. Pellets were then frozen at −80°C until needed. Pellets were thawed and resuspended in a buffer composed of 50 mM NaH_2_PO_4_, 0.5 M NaCl, and 2 mM Imidazole (pH 5) with one protease inhibitor tablet (Pierce A32965). Cells were lysed by sonication in an ice bath for 3 minutes at 60% amplitude (5 seconds pulse on; 5 seconds pulse off). The cell lysate then underwent clarification by centrifugation at 15,000 rpm (27,216 rcf) at 4°C for 15 minutes. The supernatant of AtBAM1 was then loaded onto a GE Healthcare nickel affinity column. AtBAM1 was eluted from the column using a stepwise addition of a second buffer containing 50mM NaH_2_PO_4_, 0.5 M NaCl, and 200 mM Imidazole (pH 8). The four 10 mL elution steps contained 12.5, 50, 125, or 200 mM imidazole mixed by the ÄKTA Start system. The purity of the fractions collected were then analyzed using Bis-Tris PAGE gel electrophoresis in a Tris-MOPS-SDS running buffer containing 50mM Tris Base, 50 mM MOPS, 3.5 mM SDS, and 1 mM EDTA. PageRuler Prestained Protein Ladder was used as a marker for protein size. The purest fractions as determined by SDS-PAGE were then selected and concentrated in a Spin-X UF concentrator with a PES filter that had a molecular weight cut off of 5 kDa for 30 minute intervals at 4,200 rpm (3,215 rcf) until the desired volume of 1 mL was reached. Concentrated AtBAM1 underwent further purification using the HiLoad 16/600 Superdex 200 pg on the ÄKTA Start system (Cytiva Life Science, Marlborough, MA). The protein was separated in a buffer containing 10 mM MES and 250 mM KCl (pH 7). Once the pure protein was eluted, the purity of the fractions collected were then analyzed using Bis-Tris PAGE gel electrophoresis as previously described. The purest fractions as observed by SDS-PAGE were selected and concentrated as before until the desired concentration was reached (~60 μM). The concentration of the protein was determined using Beer’s Law. The absorbance of the concentrated protein was taken at 280 nm using Synergy H4 Hybrid Reader (BioTek) on a Take3 plate, the pathlength was 0.05 cm and the extinction coefficient of 59,511 *M*^−1^cm^−1^ was calculated from the sequence using ProtParam (Gasteiger *et al.*, 2005). Concentrated AtBAM1 was flash frozen using liquid nitrogen in 50 μL aliquots, and stored at −80 °C until needed.

IbBAM5 from Sigma-Aldrich was resuspended to 7 mg/mL in 20 mM HEPES (pH 7.3), 150 mM NaCl, and 0.2 mM TCEP and separated by HiLoad 16/60 Superdex 200 column equilibrated with SEC buffer (20 mM HEPES, pH 7.3, 150 mM NaCl, and 0.2 mM TCEP).

### Size Exclusion Chromatography (SEC)

Concentrated ZmBAM7-S was further purified using a HiLoad 16/60 Superdex 200 column (Cytiva) equilibrated with SEC buffer (50 mM HEPES, pH 7.5, 25 mM NaCl, and 0.2 mM TCEP). Pure ZmBAM7-S, confirmed by SDS-PAGE, was concentrated as before, distributed into the plate for SAXS, and then flash frozen in liquid nitrogen. The concentration of ZmBAM7-S was determined using both the Bio-Rad Protein Assay Kit with BSA as the standard and by the absorbance at 280 nm using an extinction coefficient of 101,760 M^−1^cm^−1^. This value was calculated from the recombinant protein sequence including the His-tag using ProtParam *(Gasteiger et al., 2005)*.

The Size Exclusion Chromatography data for full-length and degraded ZmBAM7-S were used to predict their respective molecular weights and quaternary structures. The predicted molecular weight of a ZmBAM7 monomer was calculated using ProtParam (Gasteiger *et al.*, 2005). Experimental molecular weights were calculated using calibration standards from the Gel Filtration Molecular Weight Markers Kit for molecular weights 12-200 kDa (Sigma) and the Gel Filtration Calibration Kit for molecular weights 43-669 kDa (Cytiva). These standards were run on the HiLoad 16/60 Superdex 200 column to determine their respective void and elution volumes. The equation of the calibration curve used for molecular weight calculations was *y* =− 3. 62*x* + 10. 37using a void volume of 40.93 mL. The data were analyzed using Microsoft Excel (version 16.46), R using the tidyverse package (version 4.0.3; 10/10/2020) (Wickham *et al.*, 2019), and RStudio (version 1.3.1093) (R Core Team).

### ZmBAM7-S Homology Model Construction

Two different homology models of a ZmBAM7-S monomer were created using IntFOLD and trRosetta (Yang *et al.*, 2020; McGuffin *et al.*, 2019) from the recombinant ZmBAM7-S sequence including the N-terminal His-tag. IntFOLD selected the PDB files 1FA2 (sweet potato BAM5) and 1WDP (soybean BAM5) to use as templates for the homology model (Cheong *et al.*, 1995; Adachi *et al.*, 1998). In addition to these two templates, trRosetta also used 2XFR (barley BAM5), 1B1Y (barley BAM5), and 1BTC (soybean BAM5) as templates to create the homology model (Mikami *et al.*, 1999; Rejzek *et al.*, 2011). A DISOclust Disorder Prediction and IUPred2A were used to analyze the probability of disorder of the ZmBAM7-S protein sequence with a p-value cutoff of 0.5 (McGuffin, 2008; Mészáros *et al.*, 2018; Erdős & Dosztányi, 2020). Tetramer construction using both the IntFOLD and trRosetta homology models was completed in YASARA (version 19.5.23) by first oligomerizing the 1FA2 structure then aligning four IntFOLD-predicted or trRosetta-predicted ZmBAM7-S monomers with each 1FA2 monomer (Cheong *et al.*, 1995). Tetramer construction by alignment was followed by energy minimization using an AMBER14 forcefield with an interaction cutoff value of 12.0 Å (Maier *et al.*, 2015). To create a predicted model of truncated ZmBAM7-S for model-dependent analysis, amino acids 1-83 were deleted from the IntFOLD homology model. This truncated ZmBAM7-S model was then oligermized into a tetramer as previously described.

### Small-Angle X-ray Scattering data collection

Full length ZmBAM7-S was prepared for SAXS by diluting the SEC-purified protein using SEC buffer to five different concentrations (1.76, 2.64, 3.53, 5.29, and 6.17 mg mL^−1^) in 35 μL. A truncated form of ZmBAM7-S that appeared during purification was also prepared for SAXS by dilution in the SEC buffer; only one concentration was included (2.39 mg mL^−1^). Samples in a 96-well sample plate were flash frozen with liquid nitrogen. Three protein-free controls consisting of SEC buffer alone were included with the samples of both proteins. Purified IbBAM5 was diluted to concentrations between 1 and 10 mg mL^−1^ in 20 mM HEPES, pH 7.3, 150 mM NaCl, and 0.2 mM TCEP and flash frozen in the plate using liquid nitrogen. The sample plate was shipped overnight on dry ice to the Advanced Light Source at Lawrence Berkeley National Laboratory. Prior to data collection (date of collection: 12/07/2020), the plate was spun at 3700 rev min^−1^ for 10 minutes by beamline staff. Scattering data on the samples and controls were collected every 0.3 seconds for a total of 10 seconds resulting in 33 frames of data per sample. The beam energy was 11 keV, and the detector was 2 meters from the sample holder. Samples were kept at 10 °C during collection. Data from the protein free buffer was collected before and after ZmBAM7-S and truncated ZmBAM7-S sample collection to ensure there was no difference in scattering due to contamination of the sample cell.

For AtBAM1, data were collected on 7/22/2020 via size exclusion chromatography coupled to small angle X-ray data collection. Samples were shipped overnight at 4 °C to the SIBYLS beamline at the Advanced Light Source. The sample buffer was 50 mM MES, pH 6.7, 100 mM NaCl, 1 mM DTT. The sample was injected into an Agilent 1260 series HPLC with a Shodex KW-802.5 analytical column at a flow rate of 0.5 ml/min. Small-angle X-ray scattering (SAXS) data was collected on the eluant as it came off of the column. The incident light wavelength was 1.03 Å at a sample to detector distance of 1.5 m.

Buffer scattering was subtracted from sample scattering in RAW (version 2.0.3) using SAXS FrameSlice (version 1.4.13) as a guide to determine which frames were used during buffer subtraction (Hopkins *et al.*, 2017). This setup results in scattering vectors, q, ranging from 0.013 Å-1 to 0.5 Å-1, where the scattering vector is defined as q = 4πsinθ/λ and 2θ is the measured scattering angle.

### BAM Solution Structure Comparison

We calculated the radius of gyration and molecular weight from SAXS data for ZmBAM7-S, the truncated form of ZmBAM7-S, our previous SAXS data on AtBAM2, the reference data for sweet potato BAM from the SASBDB entry SASDA62, and data collected on sweet potato BAM from Sigma and AtBAM1 using RAW (version 2.0.3). Molecular weights were determined through Bayesian Inference. The Rg value was determined using the Guinier fit function of RAW (version 2.0.3). We then plotted the Paired-Distance Distribution function for comparison of shape and size of all the proteins. The homology model of ZmBAM7-S was fitted to the intensity data using FOXS (Schneidman-Duhovny *et al.*, 2016). All SAXS data sets have been deposited in the SASBDB as (pending codes).

### Enzyme Assays

Amylase activity assays were conducted using crude protein samples obtained by *E. coli* expression, cell lysis, and centrifugation as previously described. The sonicated supernatants of AtBAM2, ZmBAM7-S, and BL21 cells (control) were tested for amylase activity. Enzyme assays were conducted in 0.5 mL containing 50 mM 2-(N-morpholino)ethanesulfonic acid (pH 6) and various concentrations of soluble starch (Acros Organics, No. AC424495000, Morris Plains, NJ). The assays also included 100 mM KCl. After 20 minutes at 25 °C, reaction tubes were immersed in boiling water for 3 minutes to stop the reactions. The reducing sugars in each reaction were then measured using the Somogyi-Nelson assay (Nelson, 1944). Data were analyzed using Microsoft Excel (version 16.46).

## Results

Although they are evolutionarily related, BAM2 and BAM7 from *Arabidopsis thaliana* (At) are functionally and structurally quite different; AtBAM2 is a catalytically-active, plastid localized tetramer while AtBAM7 is a catalytically-inactive, nuclear localized transcription factor that probably functions as a dimer (Reinhold *et al.*, 2011; Soyk *et al.*, 2014; Monroe *et al.*, 2017, 2018; Chandrasekharan *et al.*, 2020). Interestingly, some land plants such as corn (*Zea mays;* Zm) do not have an annotated *BAM2* gene, but they contain a putative *BAM7* gene that appears to share more conserved active site residues with AtBAM2 than it does with AtBAM7. Importantly, all of the genomes that contain this unusual *BAM7* gene, which we will refer to as dual-function *BAM7* or DF-*BAM7*, also appear to lack a separate, Arabidopsis-like BAM2 gene as described by Monroe et al. (2018). Therefore, we wanted to determine if this interesting *BAM7* gene in plants that lack a *BAM2* gene encodes a structural and functional ortholog of AtBAM2 as well as AtBAM7 within the same gene. If this is true, the predicted ZmBAM2-like protein, which we hypothesize is encoded within the dual-function Zm*BAM7* gene, is initiated from a predicted second transcriptional start site (TSS), and should have catalytic and structural characteristics similar to that of AtBAM2.

### BAM2 and BAM7 Functional Residue Alignment

Arabidopsis BAM7 (AtBAM7) has an N-terminal, BZR1-like DNA-binding domain that contains a bipartite Nuclear Localization Signal (NSL) (Reinhold *et al.*, 2011). Although AtBAM7 is reported to be catalytically inactive, the BAM domain of AtBAM7 is necessary for specific DNA binding (Soyk *et al.*, 2014). The “active site” contains four mutations among the 15 residues that form H-bonds to the four glucose residues at the non-reducing end of starch (Figure 1A). These mutations likely contribute to why AtBAM7 was found to be catalytically inactive under certain conditions (Reinhold *et al.*, 2011). In the course of analyzing the BAM proteins from sequenced plant genomes, we noticed that predicted *BAM7* genes from several plants contained BAM domains that seemed to be more similar to that of AtBAM2 than to AtBAM7. We identified 12 *BAM7* genes from basal angiosperms, monocots, and basal eudicots that lacked a *BAM2* gene and compared them with the BAM2 and BAM7 protein sequences from 14 eudicot genomes that contain separate *BAM2* and *BAM7* genes. We also included BAM1 proteins from the same genomes in our analysis. We used Clustal Omega (Madeira *et al.*, 2019) to align the amino acid sequences, and then we identified residues playing a role in the specific functions of either BAM7 (Reinhold *et al.*, 2011; Soyk *et al.*, 2014) or BAM2 (Monroe *et al.*, 2014, 2018) (Figure 1A). A putative bipartite NLS was found in the BZR1-like domain of each *BAM7* gene with only minor differences. In five of the DF-BAM7 proteins the distance between the two regions of positively charged residues was 13 or 15 residues as opposed to 14 in all of the BAM7 proteins, and in half of the BAM7 and DF-BAM7 proteins, His (H) in the second positive region was substituted with Glu (E) (Figure 1A). In addition, a Glu residue that was confirmed to be essential for DNA binding (E87 in AtBAM7; (Soyk *et al.*, 2014) was perfectly conserved in all BAM7 and DF-BAM7 proteins (data not shown).

**Figure 1.**
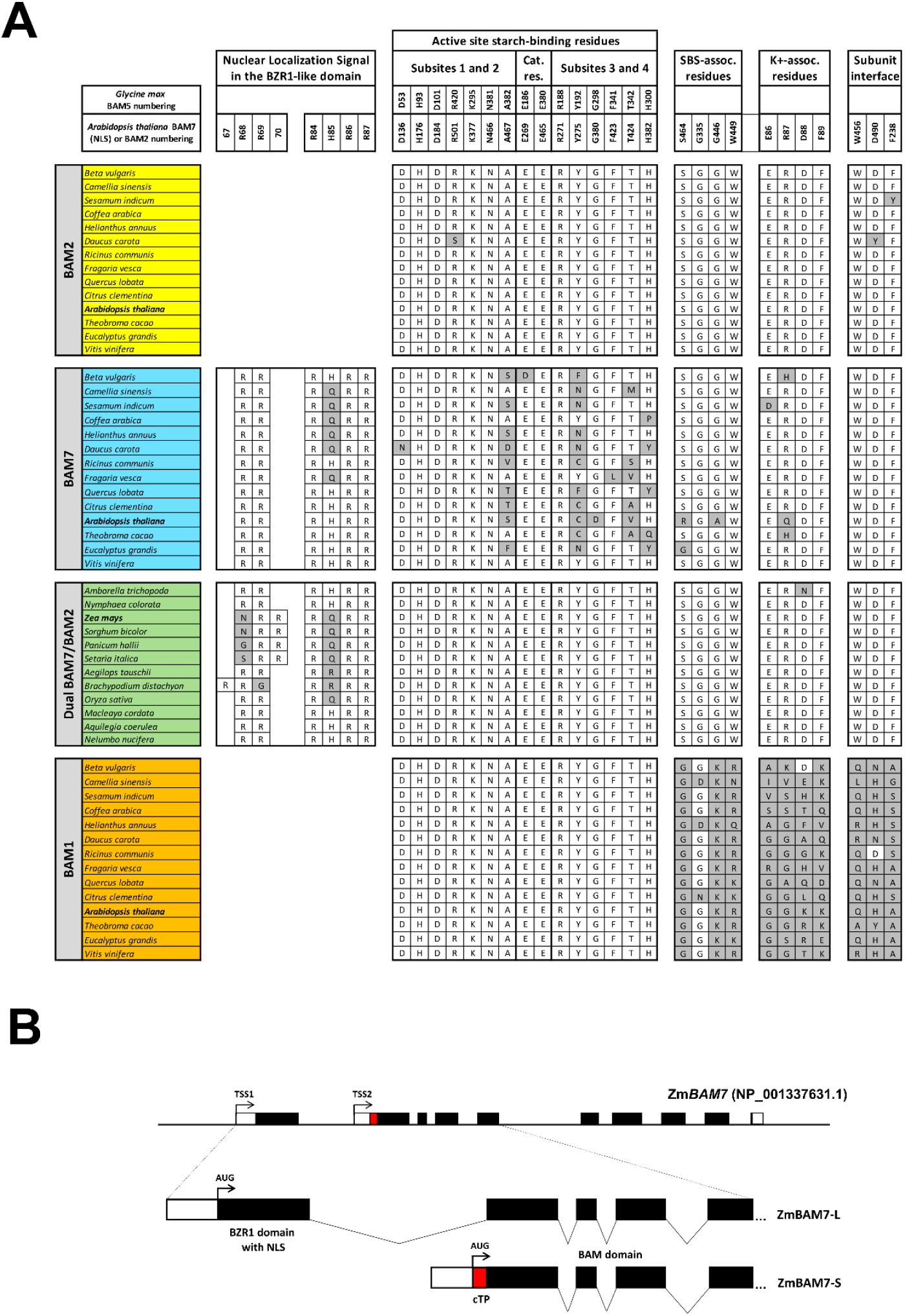
Alignment of residues known to be important for localization of BAM7, or for catalytic activity, sigmoidal kinetics, K^+^ sensitivity, or oligomerization of BAM2 as determined by Reinhold et al. (2011) and Monroe et al. (2017 and 2018) (Reinhold *et al.*, 2011; Monroe *et al.*, 2017, 2018). The yellow, blue and orange blocks of species include BAM2, BAM7 and BAM1 sequences, respectively, from 14 species that contain separate BAM2 and BAM7 genes. The green block contains 12 species in which there is a putative dual-function BAM7 gene and no BAM2 gene. Residues that differ from the consensus are highlighted with a grey background. (B) Predicted dual-function ZmBAM7 gene model. Coding regions of exons are colored black with the exception of a region unique to the N-terminus of ZmBAM7-S that is colored red. The locations of the two putative transcriptional start sites (TSS1 & TSS2) and their respective translational start sites (AUG) are indicated with arrows. The 5’ and 3’ untranslated regions (UTR) of both transcripts are colored white. Approximate locations of the nuclear localization signal (NLS) and putative chloroplast transit peptide (cTP) are labeled.

The active-site starch-binding residues (Laederach *et al.*, 1999) are perfectly conserved in all but one of the BAM2 proteins and all of the BAM1 and DF-BAM7 proteins, suggesting that these enzymes are all likely to be catalytically active. In contrast, all but one of the BAM7 sequences from eudicots that also contained a separate BAM2 gene had mutations among the active site residues (Figure 1A).

AtBAM2 is unusual among characterized BAMs in having a sigmoidal substrate saturation curve, being tetrameric, and having a secondary starch-binding site (SBS) in a groove between monomers of each dimer (Monroe *et al.*, 2017, 2018; Chandrasekharan *et al.*, 2020). We next looked for key amino acids within the BAM7 and DF-BAM7 sequences that were previously identified as functioning in each of these unique aspects of BAM2. Residues S464, E335, G446 and W449, which were previously identified as being associated with the SBS and sigmoidal kinetics in BAM2, (Monroe *et al.*, 2017, 2018) are all perfectly conserved in the BAM2 proteins and in 10 of the 12 DF-BAM7 proteins. Similarly, a 4-residue peptide (ERDF) at the N-terminus of the BAM domain that plays a role in multimerization and/or K^+^ sensitivity (Monroe *et al.*, 2018; Chandrasekharan *et al.*, 2020) was also perfectly conserved in all of the BAM2 proteins and all but one of the DF-BAM7 proteins. These residues are not conserved in the BAM1 proteins and are less well conserved in the BAM7 proteins suggesting that conservation at these positions is a hallmark of BAM2 and DF-BAM7 proteins (Figure 1A). Residues that when mutated altered oligomerization of BAM2 include F238, W456, and D490 (Monroe *et al.*, 2018). With the exception of two BAM2 proteins in which one of these residues differed, they were conserved in BAM2, BAM7 and DF-BAM7 proteins and were not conserved in the monomeric BAM1 (Sparla *et al.*, 2006). Together, these results suggest that the DF-BAM7 proteins share most, if not all, of the key residues identified as being important for BAM7 and BAM2, and thus we hypothesize that they may serve both functions.

### Dual-Function *BAM7* Gene Structure

Based on the above sequence analysis (Figure 1A), we hypothesize that the *BAM7* genes in corn and other land plants that lack a separate *BAM2* gene have alternative transcription start sites (TSSs) so that the first start site leads to a longer transcript that encodes a BAM7-like protein and contains a NLS. The shorter transcript would be initiated at the second TSS and encode a BAM2-like protein with an N-terminal chloroplast transit peptide (cTP). Using the annotated corn *BAM7* gene, we created a proposed gene structure model showing both TSSs and their respective translational start sites (Figure 1B). The longer transcript (BAM7-L) includes the first exon of the coding region which contains the DNA-binding domain and predicted NLS. This gene product would be transcribed from the first TSS, have all ten exons, and function as BAM7. The shorter transcript lacks the first exon, and would encode BAM7-S, which is initiated from a cryptic start codon at the N-terminus of a 66 amino acid-long putative cTP (Figure 1B). RNA encoding the N-terminal 14 amino acids of this cTP would be spliced out of the BAM7-L transcript, so it is specific to the BAM7-S transcript. Both transcripts have a common BAM domain with nine exons and a common translational termination site (Figure 1B). Upon closer inspection of the 12 DF-*BAM7* genes that we identified in Figure 1A, an in-frame start codon was identified in a similar position within the first intron of each gene. This cryptic translational start site is also predicted to be a part of the cTP. Therefore, the shorter versions of the DF-*BAM7* genes might be expressed and targeted to the chloroplast if they were translated from a shorter mRNA transcript. In contrast, only four of the 14 *BAM7* genes from genomes that also contained a separate *BAM2* gene had an in-frame ATG codon in exon one, and none of these four was predicted to initiate a cTP (data not shown).

### Recombinant Protein Purification

To test the hypothesis that the *BAM7* gene in some plants that lack a *BAM2* gene encodes both BAM7- and BAM2-like proteins, we expressed and purified ZmBAM7-S and AtBAM2 in *E coli*. Both ZmBAM7-S and AtBAM2 were expected to be 58 kDa including the His tag. The absence of a prominent 58 kDa band in the sonicated supernatant of BL21 cells was used to determine the expression and solubility of the recombinant proteins (Figure 2A). ZmBAM7-S and AtBAM2 were expressed (Figure 2A) and found to be soluble after sonication and centrifugation based on the 58 kD band in the sonicated supernatants of ZmBAM7-S and AtBAM2 (Figure 2A).

**Figure 2.**
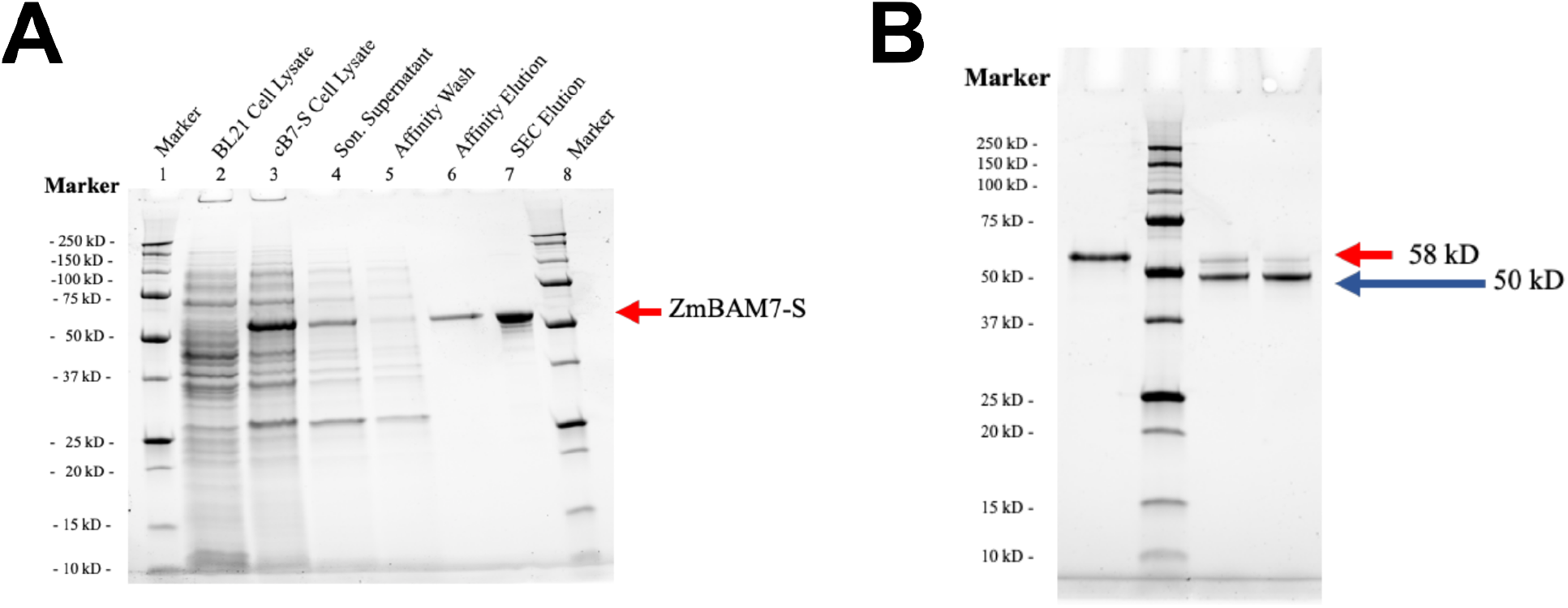
Purification of recombinant ZmBAM7-S. (A) Elutions of pure protein occurred in 10 mL fractions at increasing imidazole concentration during affinity chromatography; only one elution fraction is shown for ZmBAM7-S (lane 6). A wash fraction with less than 12.5 mM imidazole is also shown (lane 5). Size Exclusion Chromatography (SEC) was also done in preparation for Small Angle X-Ray Scattering analysis (lane 7). Markers are present in lanes 1 and 8. (B) SDS-PAGE was used to analyze an additional SEC purification of ZmBAM7-S. Prior to SEC, ZmBAM7-S was purified using an affinity column (lane 1). Two SEC elution fractions are shown (lanes 3 and 4). A protein size marker (in kDa) is in lane 2, and red and blue arrows indicate the sizes of the two bands seen in the SEC elution fractions.

We then purified ZmBAM7-S using TALON cobalt affinity chromatography followed by Size Exclusion Chromatography (SEC) (Figure 2A). The pure protein was about 58 kDa on a SDS-PAGE and the presence of multiple bands in the SEC elution indicated that ZmBAM7-S had degraded after SEC (Figure 2A). We noticed that sometimes when ZmBAM7-S was purified, degradation was not observed in the whole sample (Figure 2A), but in some cases the degradation occurred in the entire sample (Figure 2B). In these latter degradation events, the SEC elution fractions contained a minor band around the initial size of the protein (58 kDa) and a major band around 50 kDa (Figure 2B).

Following SEC, we calculated the molecular weight of ZmBAM7-S and the protein that had degraded during purification based on the elution from the column. Using these SEC elution volumes and the trend-line equation of the calibration curve, we calculated the molecular weight of truncated ZmBAM7-S to be 232.5 kDa and that of full-length ZmBAM7-S to be 384.8 kDa (Figures 3A and 3B). The molecular weight of ZmBAM7-S from the sequence should be 63 kDa suggesting that ZmBAM7-S forms oligomers of at least 4 subunits and possibly up to 6 subunits in solution.

**Figure 3.**
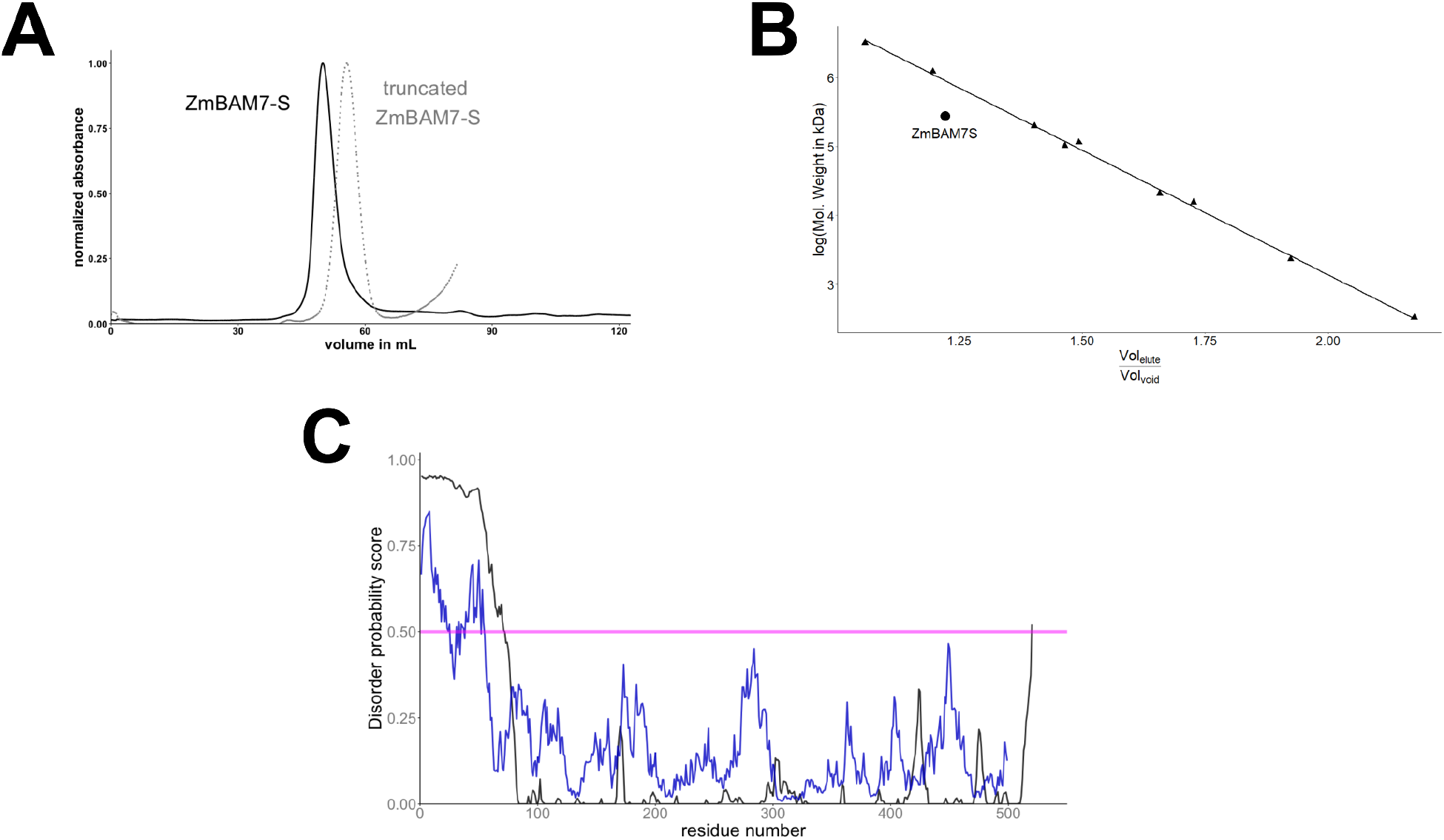
Size and disorder of ZmBAM7-S. (A) Size Exclusion Chromatogram of ZmBAM7-S and degraded ZmBAM7-S. Absorbance data were normalized to the largest value in each experiment. The peak elution volume for ZmBAM7-S (solid, black) was 50.0 mL and the peak elution volume for truncated ZmBAM7-S (dashed, grey) was 55.7 mL. (B) Size Exclusion Chromatography molecular weight calibration curve. The black line with the equation y=−3.62x+10.37 was created from nine different calibration standards (gray points). The expected ZmBAM7-S tetrameric molecular weight (black) calculated from the ZmBAM7-S sequence (232 kD) is shown. (C) Disorder predictions from IntFOLD and IUPred2A for ZmBAM7-S. Using a cutoff disorder probability score of 0.5 (pink line), the probability of being disordered was predicted for each residue in the ZmBAM7-S homology model (black line) or from the sequence of ZmBAM7-S using IUPred (blue line).

### ZmBAM7-S Homology Model

Since there are no experimentally determined structures of ZmBAM7-S, we produced homology models of the recombinant ZmBAM7-S protein sequence using IntFOLD and trRosetta (McGuffin *et al.*, 2019; Yang *et al.*, 2020). The models produced by both programs were similar except in the N-terminus since the homology model from trRosetta was more compact than that produced by IntFOLD. We then predicted the disorder of ZmBAM7-S finding that residues 1-90 of the model had the highest probability of being disordered (Figure 3C) (McGuffin, 2008; Mészáros *et al.*, 2018; Erdős & Dosztányi, 2020). The mass of the first 90 amino acids is approximately 9 kDaa consistent with the change in migration observed on SDS-PAGE between the intact and degraded proteins (Figure 2B). This finding indicates why there were inconsistencies in the two homology models and also suggests that the smaller size of truncated ZmBAM7-S may be due to loss of N-terminal residues.

### Small Angle X-Ray Scattering

We next acquired Small Angle X-Ray Scattering (SAXS) data on ZmBAM7-S and the degraded ZmBAM7-S using the SIBYLS beamline. SAXS data collection parameters and analysis software used are outlined in Table 1. We did not observe a trend in the radius of gyration (R_g_) for full-length ZmBAM7-S with concentration indicating that ZmBAM7-S forms a consistent oligomer (Figure 4A). When we compared the full-length data to that collected on the truncated ZmBAM7-S we noticed that both R_g_ and D_max_ were smaller for the truncated protein, consistent with our size exclusion data (Table 1). Kratky plots of ZmBAM7-S and the truncated protein were overall similar, suggesting that truncation does not affect the overall shape of the protein (Figure 4B). Fitting of the homology model of ZmBAM7-S to the SAXS data in FOXS showed a general match between the data and predicted data for the model, however the ^2^ value of 3.66 suggests that the tetrameric model does not fully reflect the structure (Figure 4C).

**Table 1.**

| **Sample name** | **Full Length**<br>**ZmBAM7-S** | **Truncated**<br>**ZmBAM7-S** | **AtBAM1** | **IpBAM** |
| --- | --- | --- | --- | --- |
| *Instrument* | SIBYLS beamline 12.3.1 | SIBYLS beamline 12.3.1 | SIBYLS beamline 12.3.1 | SIBYLS beamline 12.3.1 |
| *Wavelength (Å)* | 1.27 | 1.27 | 1.03 | 1.27 |
| *q range (Å<sup>-1</sup>)* | 0-0.5 | 0-0.5 | 0-0.5 | 0-0.5 |
| *Concentration (mg mL<sup>-1</sup>)* | 1.76 | 2.39 | 5.0 | 7.0 |
| *Temperature (K)* | 283.15 | 283.15 | 283.15 | 283.15 |
| *I(0) (A.U.) [from p (r)]* | 87.1 +/- 0.29 | 56.9 +/- 0.21 | 16.23 +/- 0.056 | 272.5 +/- 0.351 |
| *Rg (Å) [from p (r)]* | 51.6 +/- 0.11 | 47.4 +/- 0.13 | 27.51 +/- 0.12 | 43.67 +/- 0.051 |
| *I(0) (A.U.) [from Guinier]* | 88.9 +/- 0.7 | 61.0 +/- 0.6 | 15.68 +/- 0.075 | 272.5 +/- 0.632 |
| *Rg (Å) [from Guinier]* | 52.4 +/- 0.5 | 50.3 +/- 0.5 | 25.75 +/- 0.19 | 43.75 +/- 0.127 |
| *Dmax (Å)* | 163 | 139 | 95 | 141 |
| *Porod volume estimate (Å<sup>3</sup>) (Vp)* | 4.12x10<sup>5</sup> | 3.75x10<sup>5</sup> | 6.76x10<sup>4</sup> | 2.51x10<sup>5</sup> |
| *Molecular mass Bayes (Da)* | 318400; 100% C.I.<br>[221100-372700] | 318400; 100% C.I.<br>[221100-372700] | 53100; 100% C.I. [50300-56200] | 185800; 100% C.I.<br>[162700 – 194900] |
| *Molecular mass Mr [from Porod volume] (Da)* | 343800 | 313200 | 56100 | 208400 |
| *Calculated monomeric Mr from sequence (Da)* | 58500 | ND | 59511 | 55949 |
| *Primary data reduction* | RAW | RAW | RAW | RAW |
| *Data processing* | RAW | RAW | RAW | RAW |
| *Computation of model intensities* | FoXS | FoXS | Not Used | Not Used |
| *Three- dimensional graphic representations* | YASARA | YASARA | Not Used | Not Used |
| *SASBDB identifier* | pending | pending | pending | pending |

**Figure 4.**
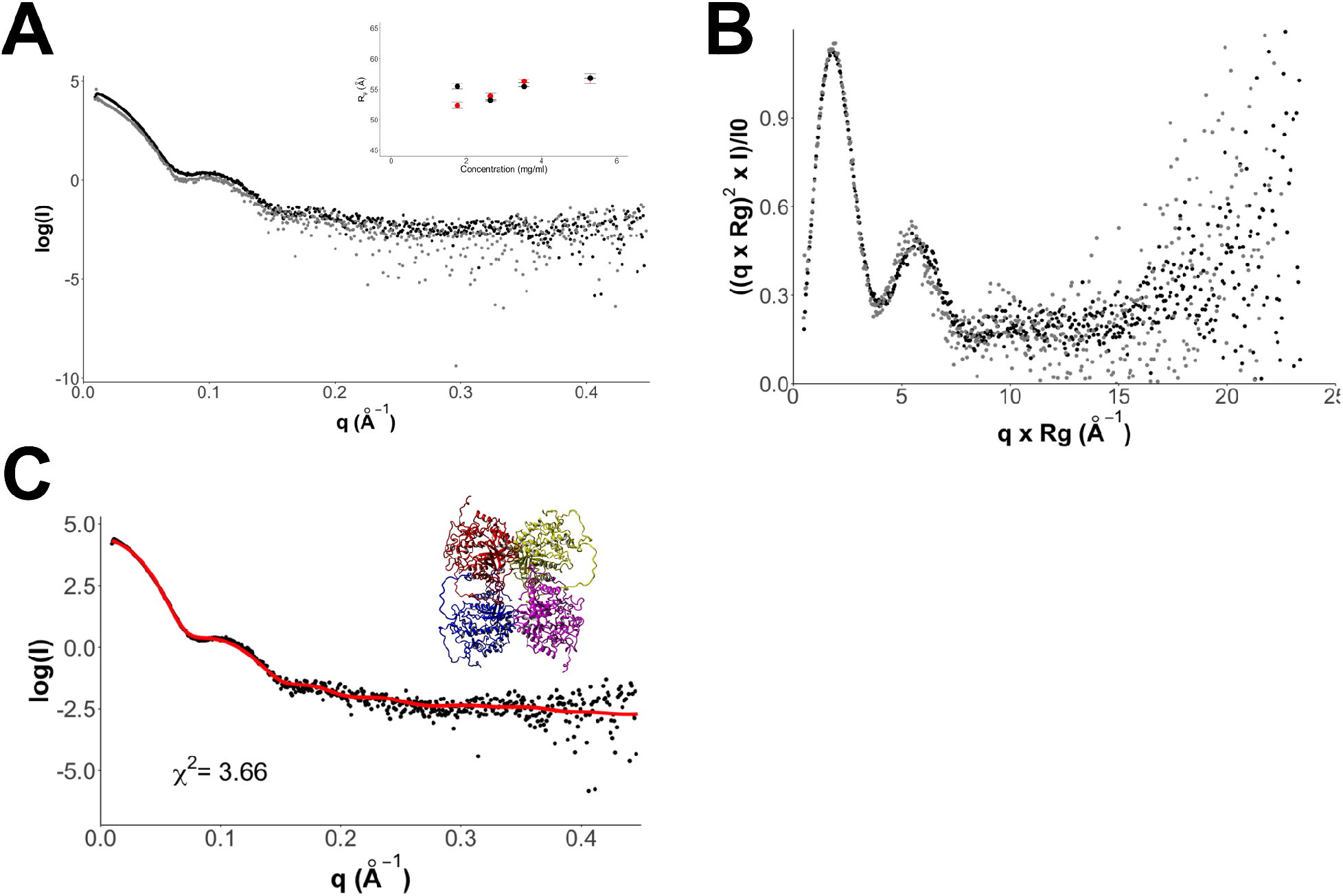
SAXS data for ZmBAM7-S and truncated ZmBAM7-S. (A) Log of intensity versus momentum transfer for ZmBAM7-S (black) and truncated ZmBAM7-S (grey) The inset plot shows radius of gyration (Rg) vs. ZmBAM7-S concentration (mg mL^−1^). The Rg value of ZmBAM-S at four different concentrations submitted for SAXS analysis were calculated from the Guinier (red data points) and from the P(r) plot (black data points). (B) Kratky plot of ZmBAM7-S (black) and truncated ZmBAM7-S (grey). (C) FOXS fitting of tetrameric homology model of ZmBAM7-S to SAXS data. The ^2^ value of the fit is 3.66. Inset shows the cartoon rendering of the tetramer used in FOXS fitting.

Because there is limited solution structure information on any BAM, we further collected SAXS data on Arabidopsis BAM1 (AtBAM1) and Ipoema BAM5 (IbBAM5) to compare to the ZmBAM7-S data. We calculated the Rg for AtBAM1 to be 25.8 ± 0.2 Å with a molecular weight of 53.1 kDa [95% CI] and for IbBAM5 to be 43.8 ± 0.1 Å with a molecular weight of 185.8 kDa [95% CI] (Table 1). IbBAM5 is generally accepted to be a tetramer in solution while BAM1 is thought to be monomeric and these data are consistent with those previous proposals (Cheong *et al.*, 1995; Sparla *et al.*, 2006). When we compared the Pair Distance Distribution Function (PDDF) for ZmBAM7-S to the data for AtBAM2, AtBAM1, and IbBAM5, we observed that the truncated ZmBAM7-S matched the data for AtBAM2, which is a homotetramer, and showed little shape similarity to BAM1, which we found to be monomeric under our conditions (Figure 5). The full-length data showed a similar peak distance value as AtBAM2, however there was a long tail toward D_max_ suggestive of a structurally extended region which was missing in the data for the truncated ZmBAM7-S. Collectively, these data support a core structure for ZmBAM7-S that is similar in shape and construction to that of AtBAM2 and IbBAM5.

**Figure 5.**
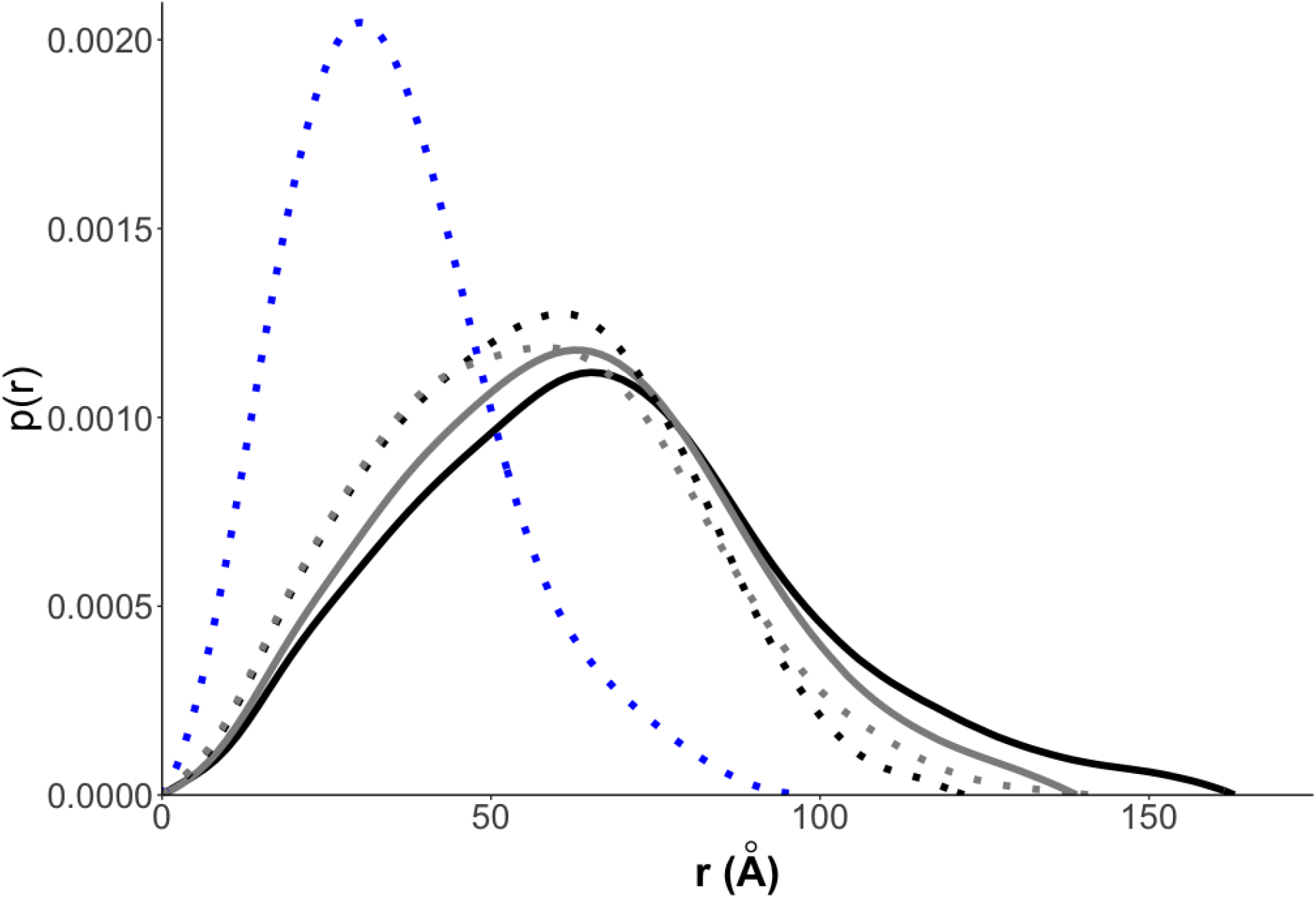
Pair distance distribution function plot comparing ZmBAM7-S and truncated ZmBAM7-S to other BAMs. ZmBAM7-S is shown as a solid black line, truncated ZmBAM7-S is shown as a solid grey line, AtBAM1 is shown as a dotted blue line, IbBAM5 is shown as a dotted grey line, and AtBAM2 is shown as a dotted black line.

### Enzyme Activity Assays

In order to avoid potentially altered activity due to the degradation of purified ZmBAM7-S (Figure 6), the sonicated supernatants of *E. coli* cells expressing ZmBAM7-S and AtBAM2 proteins were used for activity assays. After the partial purification of ZmBAM7-S and AtBAM2, the sonicated supernatants of the recombinant cells were assayed to compare their amylase activity and kinetics. The protein concentration of ZmBAM7-S and AtBAM2 could not be determined from the sonicated supernatants of the proteins because of the presence of other *E. coli* proteins in each sample. However, the proteins of interest appeared to be expressed and soluble with similar band intensities on a SDS-PAGE indicating that their concentrations in the sonicated supernatants were approximately the same. AtBAM2 is a catalytically active enzyme that degrades soluble starch *in vitro* (Monroe *et al.*, 2017). Therefore, the amylase activity of a sonicated supernatant of ZmBAM7-S was tested and compared to that of AtBAM2 with a sonicated supernatant of BL21 cells without the expression vector as a negative control (Figure 6A). The BL21 cell sample did not have any amylase activity indicating that there were no endogenous proteins in *E. coli* with BAM activity (Figure 6A inset). In contrast, both AtBAM2 and ZMBAM7-S were catalytically active, but AtBAM2 had about forty times more activity than ZmBAM7-S when assayed at 80 mg mL^−1^ soluble starch assuming similar concentrations of each protein were used (Figure 6A).

**Figure 6.**
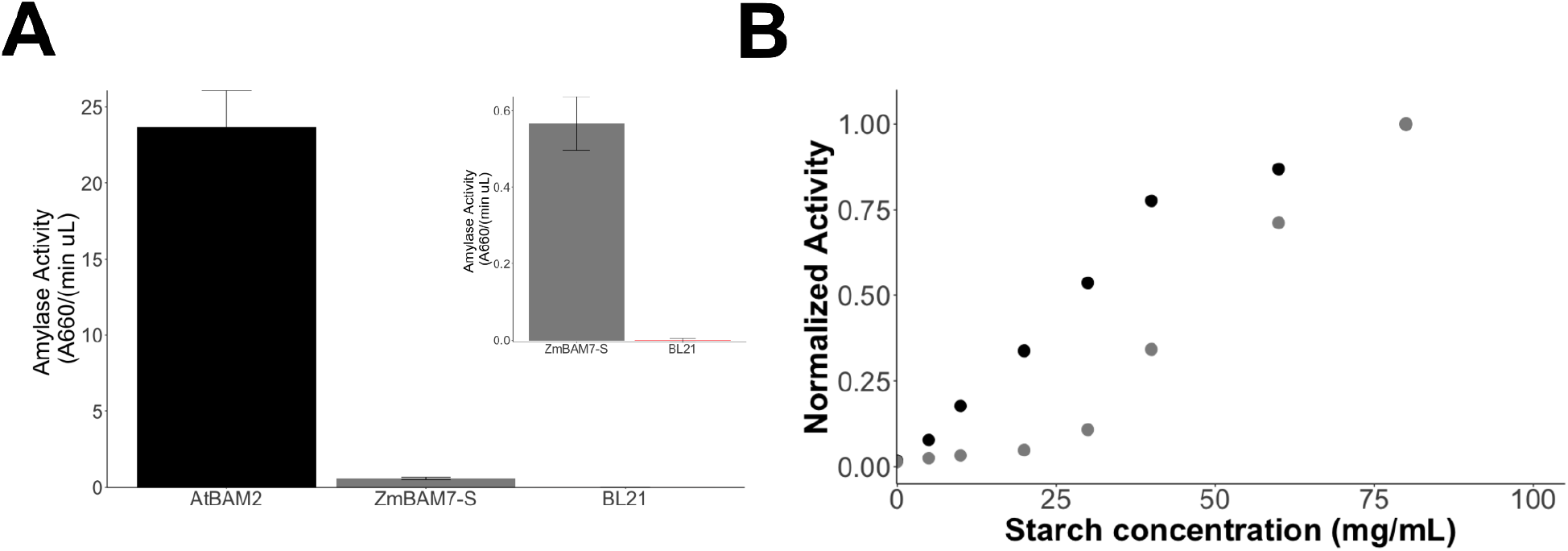
Activity of AmBAM7-S. (A) Amylase activity of the sonicated supernatant from BL21 cells (red), AtBAM2 (black), and ZmBAM7-S (grey) cells. Bars represent means +/− SD, n=3. The y-axes of both the primary graph and the internal graph both indicate amylase activity (A660 min^−1^ μL^−1^), but the y-axis of the internal graph has been expanded. Both AtBAM2 and ZmBAM7-S had similar levels of solubility as indicated by their respective bands on a gel suggesting both sonicated supernatants had comparable protein concentrations. (B) Effect of substrate concentration on ZmBAM7-S and AtBAM2 activity. Amylase activity was normalized to the activity of AtBAM2 (black) and ZmBAM7-S (grey) described in Figure 6A at the highest soluble starch concentration used (90 mg mL^−1^). Data are representative of two data sets collected.

#### Substrate Saturation

AtBAM2 is kinetically different from the other AtBAM proteins since it is the only one of those tested that exhibits sigmoidal substrate saturation kinetics (Monroe *et al.*, 2017, 2018). We performed substrate saturation experiments using AtBAM2 and ZmBAM7-S to determine if ZmBAM7-S is biochemically similar to AtBAM2 (Figure 6B). The low solubility of the soluble starch used prevented assaying ZmBAM7-S activity at a substrate concentration greater than 90 mg/mL. The amylase activity was normalized to the 90 mg mL^−1^ activity for both proteins and plotted against increasing substrate concentration (Figure 6B). The data indicate that ZmBAM7-S exhibits sigmoidal substrate binding under the assay conditions used, but the substrate binding curve of ZmBAM7-S appears to be shifted to the right of the AtBAM2 curve suggesting lower affinity (Figure 6B).

## Discussion

While analyzing the genomes of land plants for the presence of *BAM2* genes, we observed that some plants appeared to lack a *BAM2* gene but contained a *BAM7* gene. This was puzzling because *BAM2* is more ancient, and *BAM7* likely arose from a fusion of a BZR1 DNA binding domain to the 5’ end of *BAM2* (Monroe et al., 2017). However, upon closer inspection of these *BAM7* genes we discovered that the BAM domains of some have a greater percent identity to At*BAM2* than to At*BAM7*, and these *BAM7* genes have all of the catalytic and starch binding residues that *BAM7* genes sometimes lack (Figure 1A). In addition, in-frame start codons and cryptic chloroplast transit peptides were predicted within the first introns in each of the annotated *BAM7* coding regions in 12 genomes that lack a separate *BAM2* gene (Figure 1B). These observations lead us to hypothesize that the *BAM7* gene in these plants that lack a separate *BAM2* gene, such as corn, is a dual-function gene that encodes two structurally and functionally different proteins, BAM7-L and BAM7-S, by alternative transcriptional start sites, forming functional BAM7-like and BAM2-like proteins, respectively. To test this hypothesis, we designed clones of AtBAM2 and ZmBAM7-S for expression in *E. coli* to compare the catalytic and structural properties of the two proteins (Figures 6A and 6B). We also purified ZmBAM7-S using size exclusion chromatography for small angle X-Ray scattering analysis. We predicted that ZmBAM7-S would have catalytic and structural properties similar to those of AtBAM2 such as sigmoidal kinetics and a tetrameric solution structure (Figure 5).

AtBAM7 is a nuclear-localized transcription factor that likely functions as a dimer and has no previously observed catalytic activity on starch while AtBAM2 is a catalytically-active tetramer (Reinhold *et al.*, 2011; Soyk *et al.*, 2014; Monroe & Storm, 2018). If the predicted BAM7-S proteins function like AtBAM2 proteins, then they should have properties more similar to AtBAM2 than to AtBAM7. To test this, we began by analyzing the protein sequences encoded by *BAM7* genes found in some genomes that lack a separate *BAM2* gene (Figure 1A). AtBAM2’s catalytic activity has been attributed to the active site residues it shares with soybean BAM5 (1wpd); these residues include those that are necessary for starch binding and catalytic activity (Laederach *et al.*, 1999; Monroe *et al.*, 2017). In our sequence alignment, all of the putative dual-function *BAM7* genes that we analyzed have perfectly conserved active site residues like nearly all BAM2 sequences but unlike most BAM7 sequences from genomes that contain a separate *BAM2*. The BAM domain of BAM7 was found to be necessary for BAM7’s transcription factor activity, but it does not apparently catalyze a reaction like the BAM domain of BAM2 does (Soyk *et al.*, 2014). Our active site residue analysis indicated that BAM7 sequences had conservation with other BAMs in only subsite 1 and part of subsite 2. This may indicate that BAM7 binds maltose in the deep pocket of the BAM domain but does not bind starch like a BAM2-like protein does. In addition, when the full sequences of only AtBAM2, AtBAM7, and ZmBAM7 were aligned, ZmBAM7 and AtBAM7 had a conserved N-terminal sequence indicating that they potentially have similar predicted DNA-binding properties (Figure 1A). However, within the BAM domain of all three proteins, ZmBAM7 is more like AtBAM2 than AtBAM7 (Figure 1A).

In addition, ZmBAM7 shares many of the interface residues found in AtBAM2, which suggests it may form a similar tetramer. When we analyzed the SAXS data, and the PDDF derived from that data, for a full length and a fortuitously truncated form of ZmBAM7-S, we observed that the data for ZmBAM7-S aligned better to the data for AtBAM2 and IbBAM5, both of which are tetrameric (Figure 5). In comparison, the data for BAM1, which we showed was monomeric in solution, did not align with the data for ZmBAM7-S (Figure 5). The main difference between AtBAM2 and ZmBAM7-S SAXS data was the long tail in the data toward Dmax, suggesting an extended region (Figure 5). The truncated ZmBAM7-S may lack this extension, which is why the SAXS is more similar to AtBAM2 and IbBAM5. While we are not certain the exact conformation of the ZmBAM7-S structure, we are confident that the protein is tetrameric. Most BAM proteins are thought to be monomeric or a mixture of monomers and tetramers in solution, but AtBAM2 is constitutively a tetramer (Chandrasekharan *et al.*, 2020). While the physiological functions of AtBAM2 and ZmBAM7-S have not been determined, it is known that AtBAM2 is active on soluble starch and exhibits sigmoidal kinetics (Monroe *et al.*, 2017). Similarly, ZmBAM7-S was active and showed sigmoidal substrate saturation kinetics (Figures 6A and 6B). Together with the sequence and structure information, these data support our hypothesis that ZmBAM7-S is a AtBAM2-like β-amylase.

Alternative transcriptional start sites are an underappreciated mechanism of gene and protein regulation compared to alternative splicing and translational regulation. These alternative transcriptional start sites and promoters were hypothesized to regulate gene expression, alter mRNA stability, or produce two proteins with different N-terminal regions (Ayoubi & Van De Ven, 1996; Mejía-Guerra *et al.*, 2015). However others found that alternative transcriptional initiation was likely due to molecular errors and was not adaptive (Xu *et al.*, 2019). Genome-wide transcriptional start site (TSS) determination in corn identified about 1,500 genes that have multiple transcriptional start sites (Mejía-Guerra *et al.*, 2015). Sequenced cDNAs in the Maize Genome Database (https://www.maizegdb.org/) from corn (locus Zm00001d019756) appear to encode both long and short BAM7 proteins (Portwood *et al.*, 2019). In addition, our preliminary analysis of ZmBAM7 transcripts using 5’ RACE also supports that long and short transcripts of this gene exist *in vivo* (K. Ozcan and J. Monroe, unpublished data). If the corn DF-BAM7 gene indeed encodes two functionally distinct proteins orthologous to Arabidopsis BAM7 and BAM2, then it is conceivable that other unrecognized functional genes reside within annotated genomes. Techniques for identifying alternative transcriptional start sites within coding regions would be useful in correcting these oversights and preventing more in the future.

## Acknowledgements

This work was supported by a National Science Foundation Research at Undergraduate Institutions grant to JDM and CEB (MCB-1932755), a Summer Undergraduate Research Fellowship from the American Society of Plant Biologists to CMR, National Science Foundation Research Experience for Undergraduates (CHE-1757874), infrastructure provided by a grant from the Thomas F. and Kate Miller Jeffress Memorial Trust to CEB, and a grant from the 4-VA organization to CEB. Additional support comes from the National Institutes of Health project ALS-ENABLE (P30 GM124169) and a HighEnd Instrumentation Grant (S10OD018483).

