## Supplemental Information 1 for "The *β-amylase7* gene in *Zea mays* encodes a protein with structural and catalytic properties similar to Arabidopsis BAM2"

*By*

Claire M. Ravenburg, McKayla B. Riney, Jonathan D. Monroe, and Christopher E. Berndsen

Supplemental Information 1

>AtBAM2

AtgggcagcagccatcatcatcatcatcacagcagcggcctggtgccgcgcggcagcCATATGGCGGAGAGCACCGAGGAAGATCGTGTTCCGATCGACGATGACGATGACAGCACCGATCAGCTGGTTGACGAGGAAATTGTGCACTTTGAGGAACGTGACTTTGCGGGTACCGCGTGCGTGCCGGTTTACGTGATGCTGCCGCTGGGCGTGATCGATATGAACAGCGAGGTGGTTGAACCGGAGGAACTGCTGGACCAGCTGCGTACCCTGAAGAGCGTTAACGTGGATGGTGTTATGGTGGACTGTTGGTGGGGCATTGTTGAAAGCCACACCCCGCAAGTGTACAACTGGAGCGGTTATAAGAAACTGTTTCAGATGATCCGTGAGCTGGGCCTGAAAATTCAAGTGGTTATGAGCTTCCACGAATGCGGTGGCAACGTTGGTGATGACGTGCACATCCAGATTCCGGAGTGGGTTCGTGAAATCGGTCAAAGCAACCCGGATATTTATTTTACCGACAGCGCGGGCCGTCGTAACACCGAGTGCCTGACCTGGGGTATCGATAAGCAACGTGTTCTGCGTGGCCGTACCGCGCTGGAAGTGTACTTCGATTATATGCGTAGCTTTCGTGTTGAGTTCGACGAATTCTTTGAGGAAAAGATCATTCCGGAGATCGAGGTTGGTCTGGGTCCGTGCGGCGAGCTGCGTTACCCGAGCTATCCGGCGCAGTTTGGCTGGAAATACCCGGGTATTGGCGAATTCCAATGCTACGACAAGTATCTGATGAACAGCCTGAAAGAGGCGGCGGAAGTGCGTGGTCACAGCTTCTGGGGTCGTGGCCCGGATAACACCGAAACCTATAACAGCACCCCGCACGGTACCGGCTTCTTTCGTGACGGTGGCGATTACGACAGCTACTATGGTCGTTTCTTTCTGAACTGGTATAGCCGTGTTCTGATCGATCACGGCGACCGTGTGCTGGCGATGGCGAACCTGGCGTTTGAGGGTACCTGCATCGCGGCGAAGCTGAGCGGCATTCACTGGTGGTACAAAACCGCGAGCCATGCGGCGGAACTGACCGCGGGTTTCTACAACAGCAGCAACCGTGATGGTTATGGCCCGATTGCGGCGATGTTTAAGAAACACGACGCGGCGCTGAACTTCACCTGCGTTGAGCTGCGTACCCTGGATCAGCACGAGGACTTTCCGGAAGCGCTGGCGGACCCGGAAGGCCTGGTTTGGCAAGTGCTGAACGCGGCGTGGGATGCGAGCATTCCGGTGGCGAGCGAAAACGCGCTGCCGTGCTACGATCGTGAGGGTTATAACAAGATTCTGGAAAACGCGAAACCGCTGACCGATCCGGATGGTCGTCACCTGAGCTGCTTCACCTACCTGCGTCTGAACCCGACCCTGATGGAGAGCCAGAACTTCAAGGAGTTTGAACGTTTCCTGAAACGTATGCACGGTGAAGCGGTTCCGGACCTGGGTCTGGCGCCGGGTACCCAAGAAACCAACCCGGAATAA

>ZmBAM7-S

ATGGGCAGCAGCCATCATCATCATCATCACAGCAGCGGCCTGGTGCCGCGCGGCAGCCATATGGCGGCGGCGAGCCGTAGCCCGGCGGATGATGTTCCGGACGGTAACAGCAGCCACCTGCTGGCGGTTCCGGTTCCGGTGCCGATGGACCCGGCGGCGGCGGAAGATGTTCCGGTGGCGAAGCAGCTGCAAGTTCCGGATGTTAGCCCGCGTCCGCCGGAGCGTGATTTTGCGGGCACCCCGTACGTTCCGGTGTATGTTATGCTGCCGCTGGGTGTGGTTAACGGTAACGGCGAAGTGGTTGACGCGGATGAGCTGGTTGGTCAGCTGCGTGTTCTGAAGGCGAGCGGTGTGGACGGCGTGATGGTTGATTGTTGGTGGGGCAACGTTGAAGCGCACAAACCGCAAGAGTACAACTGGACCGGTTATCGTCGTCTGTTTCAGATGATCCGTGAACTGAAGCTGAAACTGCAAGTGGTTATGAGCTTCCACGAGTGCGGTGGCAACGTGGGTGACGATATCAGCATTCCGCTGCCGCACTGGGTTATCGAAATTGGCCGTAGCAACCCGGACATTTACTTTACCGATCGTGCGGGTCGTCGTAACACCGAATGCCTGAGCTGGGGCGTGGACAAAGAGCGTGTTCTGCAGGGTCGTACCGCGGTGGAAGTTTACTTCGATTTTATGCGTAGCTTCCGTGTGGAGTTTGACGAATATTTCGAGGATGGTATCATTAGCGAGATCGAAATTGGTCTGGGTGCGTGCGGTGAACTGCGTTACCCGAGCTATCCGGCGAAGCACGGTTGGAAATACCCGGGTATCGGCGAATTCCAGTGCTACGACCGTTATCTGCAAAAGAGCCTGCGTAAAGCGGCGGAGGCGCGTGGTCACACCATTTGGGCGCGTGGTCCGGACAACGCGGGTCACTATAACAGCGAGCCGAACCTGACCGGCTTCTTTTGCGATGGTGGCGACTACGATAGCTACTATGGTCGTTTCTTTCTGAGCTGGTATAGCCAGGCGCTGGTTGACCACGCGGATCGTGTTCTGATGCTGGCGCGTCTGGCGTTTGAAGGCACCAACATCGCGGTGAAAGTTAGCGGTGTGCACTGGTGGTACAAAACCGCGAGCCATGCGGCGGAGCTGACCGCGGGCTTCTACAACCCGTGCAACCGTGACGGTTATGCGCCGATTGCGGCGGTTCTGAAGAAATACGATGCGGCGCTGAACTTTACCTGCGTGGAACTGCGTACCATGGACCAACACGAAGTTTATCCGGAGGCGTTTGCGGACCCGGAGGGTCTGGTGTGGCAAGTTCTGAACGCGGCGTGGGATGCGGGTATCCAAGTTGCGAGCGAAAACGCGCTGCCGTGCTACGACCGTGATGGCTTCAACAAGATTCTGGAGAACGCGAAACCGCTGAACGACCCGGATGGCCGTCACCTGCTGGGTTTTACCTATCTGCGTCTGGGCAAGGACCTGTTCGAACGTCCGAACTTCTTTGAGTTCGAACGTTTTATCAAACGTATGCACGGTGAGGCGGTGCTGGATCTGCAAGTTTAA

**Figure S1.** Sequences of AtBAM2 and ZmBAM7-S synthesized by GenScript and cloned into pET15a for expression in *E. coli*.
